# Cyclic, condition-independent activity in primary motor cortex predicts corrective movement behavior

**DOI:** 10.1101/453746

**Authors:** Adam G. Rouse, Marc H. Schieber, Sridevi V. Sarma

## Abstract

Reaching movements are known to have large condition-independent neural activity and cyclic neural dynamics. A new precision center-out task was performed by rhesus macaques to test the hypothesis that cyclic, condition-independent neural activity in the primary motor cortex (M1) occurs not only during initial reaching movements but also during subsequent corrective movements. Corrective movements were observed to be discrete with time courses and bell-shaped speed profiles similar to the initial movements. Condition-independent cyclic neural trajectories were similar and repeated for initial and each additional corrective submovement. The phase of the cyclic condition-independent neural activity predicted the time of peak movement speed more accurately than regression of instantaneous firing rate, even when the subject made multiple corrective movements. Rather than being controlled as continuations of the initial reach, a discrete cycle of motor cortex activity encodes each corrective submovement.

**Significance Statement:** During a precision center-out task, initial and subsequent corrective movements occur as discrete submovements with bell-shaped speed profiles. A cycle of condition-independent activity in primary motor cortex neuron populations corresponds to each submovement, such that the phase of this cyclic activity predicts the time of peak speeds—both initial and corrective. These submovements accompanied by cyclic neural activity offer important clues into how we successfully execute precise, corrective reaching movements and may have implications for optimizing control of brain-computer interfaces.

## Introduction

Corrective movements based on sensorimotor feedback are critical for elegant motor control. While a single, discrete movement like a pointing gesture may be mostly ballistic, more precise aiming movements typically require an error correction phase (Woodworth, 1899; Craik, 1947; Abrams et al., 1990; Sainburg et al., 1999; Elliott et al., 2010). In making an online correction, the brain must respond to updated sensory information about the current position relative to the desired target. Yet the way neurons in motor areas of the brain encode and generate corrective movements to achieve movement precision is relatively unexplored. When examining populations of neurons in primary motor cortex during instructed movements, predictable dynamics of neural spiking occur with a progression from initiation to completion of a movement (Maynard et al., 1999; Jackson et al., 2003; Truccolo et al., 2005; Sarma et al., 2010). Yet behaving animals also respond to updated sensorimotor information, as happens in tasks that require precision. For corrective movements with new sensory information, does the neural activity update within a current active neural state as a continuation of the initial reach or does it repeat and cycle again through the same series of neural dimensions for each additional submovement?

We investigated the neural dynamics underlying corrective movements, focusing on two key features of neural activity in primary motor cortex that have been previously described during reaching: i) condition-independent neural activity and ii) rotations in neural dynamics. Although individual neurons in primary motor cortex encode a variety of condition-dependent movement features (Evarts, 1968; Thach, 1978; Georgopoulos et al., 1982; Kalaska et al., 1989; Kakei et al., 1999), there is also a large condition-independent component in the firing rate of neurons in motor cortex (Kaufman et al., 2016; Rouse and Schieber, 2018). Condition-independent neural activity is the change in a neuron’s firing rate from baseline over time that happens regardless of the instructed movement for any given trial within a given task. Condition-independent activity presumably carries information on the timing of movement as opposed to specific, condition-dependent features. Techniques like demixed principal component analysis can partition a neural population’s activity into condition-independent modulation and the more classically described condition-dependent tuning to task conditions (Kobak et al., 2016). In addition to being condition-independent or -dependent, changes in firing rate in theory might be temporally synchronous across a population. But in practice, primary motor cortex neurons have an asynchronous range of onset latencies before movement, with latencies for most corticomotoneuronal (CM) cells ranging from 120ms to 0ms (Cheney and Fetz, 1980) while other motor cortex neurons can lead movement by up to 200ms (Moran and Schwartz, 1999). Because the increases and decreases in firing rates are not synchronous, the population activity forms a more complex trajectory in neural state space (Yu et al., 2007; Cunningham and Yu, 2014). These time-varying dynamics can either be dependent on specific task conditions or independent of task conditions. While the precise meaning of these features of neural dynamics under different conditions remains debated (Churchland et al., 2012; Hall et al., 2014; Michaels et al., 2016; Lebedev et al., 2019), these shifts between different combinations of active neurons leads to changing dimensions of the neural space.

We hypothesized that if the primary motor cortex handles online corrections as ongoing adjustments to a single reach, then one cycle of the neural trajectory would include both the initial and the corrective submovements. In contrast, if the primary motor cortex handles each correction as a distinct (albeit smaller) movement, then each corrective submovement would correspond to its own cycle repeating the series of neural dimensions that are traversed. We used a precision center-out task that required moving to small targets (either narrow or shallow) to elicit visuomotor corrections. We examined whether corrective movements in this task were simple adjustments in the ongoing reach or discrete submovements, behaviorally similar to initial movements. We then ask whether condition-independent activity—representing the time course of movement irrespective of its direction or amplitude—is similar for both initial and corrective submovements. Finally, we ask whether cyclic neural dynamics improve our predictions of when initial and corrective movements occur.

## Materials and Methods

### Non-human primates

Two male rhesus monkeys, P and Q (weight 11 and 10 kg, ages 7 and 6 years old, respectively), were subjects in the present study. All procedures for the care and use of these nonhuman primates followed the Guide for the Care and Use of Laboratory Animals and were approved by the University Committee on Animal Resources at the University of Rochester Medical Center, Rochester, NY.

### Experimental Design

A precision center-out task was performed by the monkey, using an 18 cm handle attached to a commercial joystick (M212 series joystick, PQ Controls Inc.) to control a cursor on a 24” LCD display. The joystick handle moved freely with minimal resistance as the spring mechanism for providing centering, restorative force was removed. The end of the joystick could move approximately 9.3 cm in both the forward/backward and left/right directions. Motion of the joystick was transduced linearly by two Hall effect sensors sliding in both the backward/forward and left/right directions. The cursor viewed by the monkey directly represented the planar position of these two sensors scaled to fit within a 1000 horizontal x 1000 vertical pixel workspace in the center of the LCD display. The limits of the cursor workspace were slightly within the physical limits of the joystick, with 110 pixels corresponded to approximately 1 cm of movement at the end of the joystick. The cursor appeared on the display as a small cross centered on a single pixel in the workspace. Custom software for task control sampled the joystick data, updated the scene, and stored the cursor position (equivalent to joystick position) and trial event times at 100 Hz.

The precision center-out task consisted of three sets of eight peripheral targets located equidistance and equally spaced in 45° intervals around a center, home target (see Figure 2). The center target had a radius of 75 pixels. Each center-out target—defined in polar coordinates—was one of three different sizes i) large targets spanning 45° of the workspace and covering 250-450 pixels from the center, ii) shallow targets spanning 45° but covering a width of only 325-375 pixels from the center, and iii) narrow targets spanning 15° covering 250-450 pixels from the center. All 24 targets (3 sizes x 8 locations) were presented pseudo-randomly in equal amounts throughout a session.

For each trial, following the subject acquiring the home target and performing a required initial hold ranging from 300-500 ms, the instruction occurred with the given trial’s correct target changing from black to green. Following this instruction, the monkey could move the cursor immediately to contact the correct target. At contact, the outline of all targets changed colors from white to black providing visual feedback that the cursor was within the target boundaries. After contacting the desired target, the cursor was required to remain within the target for a variable hold time of 500-600 ms. If the cursor left the target during this hold, the monkey was allowed to enter the target again and complete a final hold. Once a successful final hold of 500-600 ms was completed, the animal received a liquid reward. Both the required initial and final hold times for each trial were randomly sampled from a uniform distribution.

### Neural Recordings

Floating microelectrode arrays (MicroProbes for Life Science) were implanted in the anterior lip and bank of the central sulcus to record from primary motor cortex (M1) in each monkey, using methods described in detail previously (Mollazadeh et al., 2011; Rouse and Schieber, 2016). For monkey P, recordings were collected from six 16-channel arrays implanted in M1. For monkey Q, two 32-channel arrays and one 16-channel array in M1 were used. The location of the implanted arrays, spanning the forelimb representation in M1, have been previously reported (Fig. 2 of (Liu and Schieber, 2020)) and spanned the forelimb area of M1. Intracortical microstimulation on single electrodes with a current up to a maximum of 100 μA (12 biphasic pulses, 0.2ms pulse width per phase, 3ms interpulse interval) with the animal lightly anesthetized with ketamine evoked a variety of forelimb movements. Of the 96 electrodes for monkey P, stimulation of 11 sites elicited proximal arm movements, 6 sites elicited wrist movements, and 21 sites elicited movement of the digits. Of the 80 electrodes for monkey Q, 34 sites were proximal, 9 sites were wrist, and 25 were digits. During recording sessions, channels with spiking activity were thresholded manually online, and spike-waveform snippets and spike times were collected with Plexon MAP (Plexon, Inc.) and Cerebus (Blackrock Microsystems, LLC.) data acquisition systems. The spike snippets were sorted off-line with a custom, semi-automated algorithm. Chronic multielectrode arrays do not always yield well-isolated single-unit recordings. To define likely single units, we utilized the signal to noise ratio of the sorted spike waveforms and the percent of true single unit spikes estimated from a formula using the number of interspike interval (ISI) violations less than 1ms (Hill et al., 2011; Rouse and Schieber, 2016). Using a signal to noise ratio of SNR > 3 and 100% true single unit spikes (no ISI violations) to define definite single units and SNR > 2.5 and >90% true single unit spikes to define probable single units, 543 (monkey P) and 304 (monkey Q) of sorted spike waveforms were classified as definite single units while 268 (P) and 208 (Q) additional units were probable single units. Thus, 811/1293=63% (monkey P) and 512/1185 = 43% (monkey Q) of all spiking units were classified as likely single units. Because the estimation of neural population states from multi-unit activity has previously been shown to be quite similar to that from well isolated single units (Trautmann et al., 2019) and because including multi-units would be unlikely to provide results more significant than similar numbers of single-units, we included both single- and multi-unit recordings in our analyses.

### Behavior Analyses

A peak finding algorithm to identify local maxima was used for analysis of the timing of cursor speed peaks. Off-line, cursor speed was calculated by filtering the cursor position with a 10-Hz low-pass 1^st^-order Butterworth filter (bidirectionally for zero phase lag) and then calculating the first derivative using the 5-point central difference. Local maxima of cursor speeds (identified with *findpeaks* function in Matlab (Mathworks, 2020)) were identified as peaks if they met the following criteria: i) the peak speed was greater than 250 pixels/s and ii) the peak’s prominence— the height difference between the peak and the larger of the two adjacent troughs (minimum speed before encountering a larger peak)—was at least 50% of the absolute height of the peak. All such cursor speed peaks with their surrounding ±200 ms time windows were considered submovements within a trial. Initial peaks were identified as the first submovement that ended at least 150 pixels from the center (approximately halfway to the peripheral target). Any small movements before the initial speed peak—506 (4.6% of trials) for P and 616 (7.0% of trials) for Q—were discarded from further analysis. Speed peaks following the initial speed peak were defined as corrective submovements. To focus analysis on submovements made to successfully acquire the target, corrective submovements were only included if some portion of the acceleration phase—time from preceding speed trough to speed peak—occurred outside the peripheral target.

The speed profiles for individual submovements were analyzed between -200 and 200 ms relative to peak speed. As a measure of similarity between speed profiles, the Pearson’s correlation between these speed profiles for pairs of submovements was calculated, yielding a similarity score between -1 and 1. To measure how similar corrective submovements were to initial submovements, the correlation of each initial submovement to a randomly selected corrective submovement was calculated. As a ceiling comparison, each initial submovement was also compared to another randomly selected initial submovement. Thus, the distribution of correlations for initial-corrective submovement pairs was compared to the distribution of initial-initial pairs.

### Identifying condition-independent, rotational neural activity

We focused our neural population analysis on the neural dimensions that contained the most condition-independent, rotational activity. A schematic illustration of these two features—i) condition-independent vs. -dependent, and ii) synchronous vs. rotational/asynchronous is shown in Figure 1. The condition-independent activity is the time-varying average of firing rate across all trials regardless of condition while the condition-dependent is the specific tuning to task condition like target direction. Synchronous, time-locked activity represents changes in firing rate that happen simultaneously across the neural population, while asynchronous activity of varying time course in different neurons can lead to patterns of traveling waves or oscillations in the population with a predictable progression in time.

**Figure 1.**
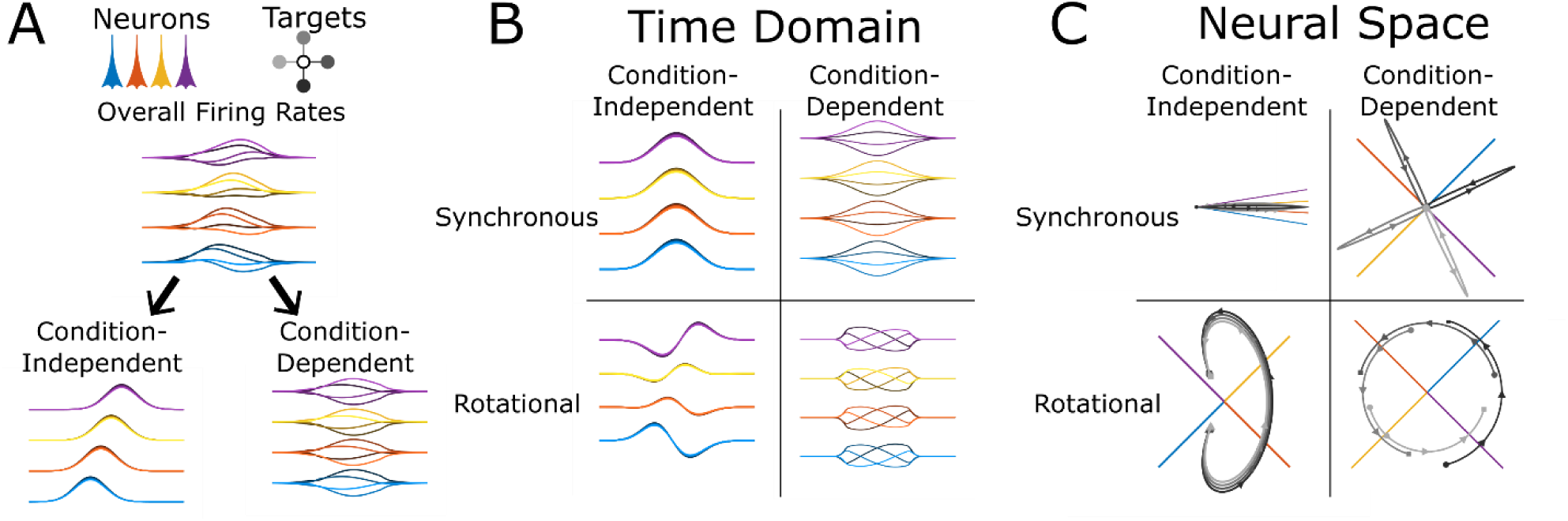
Idealized representation of both the synchronous and rotational components of condition-independent and -dependent changes in neuronal firing rate. A) The firing rates for four neurons (blue, orange, yellow, and purple) are shown for reaches to four target directions (light to dark grayscale). The overall firing rates differ for both the four neurons and the four target condition. By time averaging across the four conditions, the condition-independent firing rates and the residual condition-dependent firing rates are both identified. B) Next, averaging across the population reveals that firing rates are i) synchronous activity across all neurons at each time point and ii) the remaining, asynchronous/rotational firing rate changes specific for each neuron. C) The neural space visualizes the population activity by showing each neuron’s firing rate as a point along an orthogonal dimension with time represented as a trajectory through this space. In this representation, the difference between synchronous and rotational activity is better appreciated. Synchronous activity is movement along a single neural dimension while rotational activity is movement between dimensions. Note, the dimensions defined by individual neurons are shown projected in a 2D plane. Only the given component (synchronous/rotational and condition-independent/-dependent) are shown for these four example neurons for visualization purposes. In a much higher dimensional space when recording from a large number of neurons, the possibility of finding dimensions with little overlap between components is much greater.

Firing rates of the neural population can be visualized as either: i) a function of time (Fig. 1B) or ii) neural trajectories in a Cartesian neural space where each neuron’s firing rate is plotted on an orthogonal dimension (Fig. 1C). For a complex task with variable corrective submovements such as our precision center-out task, the condition-independent activity provides a useful analysis to identify the neural activity underlying a submovement. Although a synchronous rise and fall of firing rate across the neural population—a single neural dimension--may provide some information, utilizing additional neural dimensions of the condition-independent signal may help improve our prediction of the timing and phase of submovements. The simplest is to consider two-dimensions of condition-independent activity in which the rotational activity resulting from sequential firing rate changes across different neurons produces a cycle in a neural plane. This approach has the potential to improve identification of corrective submovements.

### Dynamical Systems Model

Traditionally, condition-independent signals are identified by aligning neural data to behavioral cues and time averaging with methods like dPCA (Kaufman et al., 2016; Ames and Churchland, 2019). However, our precision center-out task consisted of corrective movements that were highly variable in their timing relative to any experimental controlled behavioral event. We therefore employed dynamical system modeling to characterize repeated changes in firing rates across our recorded neural population. To identify and analyze potential repeatable temporal dynamics of the neural population that correlated with movement, our neural data was modeled as a linear, time-invariant system using a system of coupled first-order ordinary differential equation defined by a transform matrix. This model was built using only the condition-independent activity by averaging the firing rates for individual spiking units across all trials regardless of the movement condition (i.e. target location).

The condition-independent activity was then submitted to jPCA (Churchland et al., 2012) to identify the two-dimensional neural plane with the most rotational/cyclic activity. In this model, the changes in firing rate can grow/shrink along a single dimension (synchronous) as well as rotate across dimensions (asynchronous). The eigen decomposition of the transform matrix yields eigenvalues with the real part representing growing or shrinking away from the origin while the imaginary part represents rotations. Note, this utilization of the jPCA algorithm on only the condition-independent activity is different than the typical application of jPCA to data containing the condition-dependent activity. Additionally, we find the results of the dynamical system are more stable when the firing rates are square-root transformed to equalize variance between high and low firing rates (Kihlberg et al., 1972; Snedecor and Cochran, 1980; Ashe and Georgopoulos, 1994) and thus performed this transform before submitting firing rates to jPCA.

We call the plane with the most rotation the condition-independent (CI) plane and define the two neural dimensions that define this plane as CIx and CIy. To consistently define CIx and CIy across recording sessions and monkeys, we defined the +CIx direction as the neural dimension that had the maximum average firing rate. This was performed by calculating the population averaged firing rate at all angles in the plane and rotating the CIx and CIy axes so that +CIx aligned with the largest firing rate. Having identified this jPC neural plane, our work introduces a new analytic variable—condition-independent phase (CIφ)—which estimates the instantaneous phase angle within this two-dimensional plane of the projected population firing rates. We calculate CIφ using the Hilbert transform applied to the two signals, CIx and CIy, generating a complex, analytical representation of the population signal. The angle of this complex signal is then used to calculate the instantaneous phase.

Since our task consisted of highly variable trial lengths and timing, the identification of condition-independent activity by time averaging based upon behavioral events was challenging. To be less constrained in identifying the plane with condition-independent rotational activity, we used an iterative approach alternating between identifying the CIφ for each time point and then averaging the condition-independent neural activity for each CIφ value. We first time-averaged the activity aligned on speed peaks, and then initially performed jPCA on the time-averaged data. After identifying the rotational plane, we then binned and averaged the firing rates based on its phase in the plane (rather than time) and performed jPCA on this new phase-averaged neural activity. This calculation of the jPCA plane and phase averaging was repeated for three iterations to ensure convergence. The Matlab code and additional documentation about the calculation of CIφ as described in the paper is freely available online at https://github.com/arouseKUMC/CIphase. The code is also available as Extended Data 1.

The calculation of the jPC plane and the CIφ was performed using 5-fold cross-validation. Each recording session was divided into 5 testing sets of trials each containing 20% of the data. The jPC plane was calculated by training on the other 80% of the data and then tested on each test set. All presented results for CIφ are using the test data projected into the jPC dimensions identified by the separate training set.

### Firing Rate vs. Speed Model

For comparison with our two-dimensional CI plane and phase analysis, we wanted to examine how well a linear predictor of speed using a single neural dimension could perform. We therefore performed linear regression to predict speed from the recorded neural firing rates. For this estimate, we regressed the firing rates for all recorded units to peak speed for all submovements. We utilized the firing rates for each recorded unit averaged across a time window from 300 ms before to 100 ms after each peak speed. We chose this method to identify a neural dimension that correlated with speed without using separate time lags for each individual neuron. For motor cortex, the neural signal in this dimension would be expected to increase and peak before each peak in movement speed. We identify and report the time at which the peaks in this neural signal occurred to quantify how accurately the timing of peaks in movement speed was predicted.

### Statistics

Several statistical analyses (Table 1) were used to assess how similar corrective submovements were to initial submovements and whether there were repeated cycles of neural activity and if these cycles corresponded to behavior. For correlations between submovement speed profiles, movement times, and average spike times, non-parametric tests were used. Since CIφ values represent an angle ranging from -π to π, circular distribution statistics—mean, variance, correlation, and Rayleigh test for non-uniformity— were used. All circular statistics were calculated with CircStat, a Circular Statistics Toolbox for Matlab (Berens, 2009).

**Table 1.**
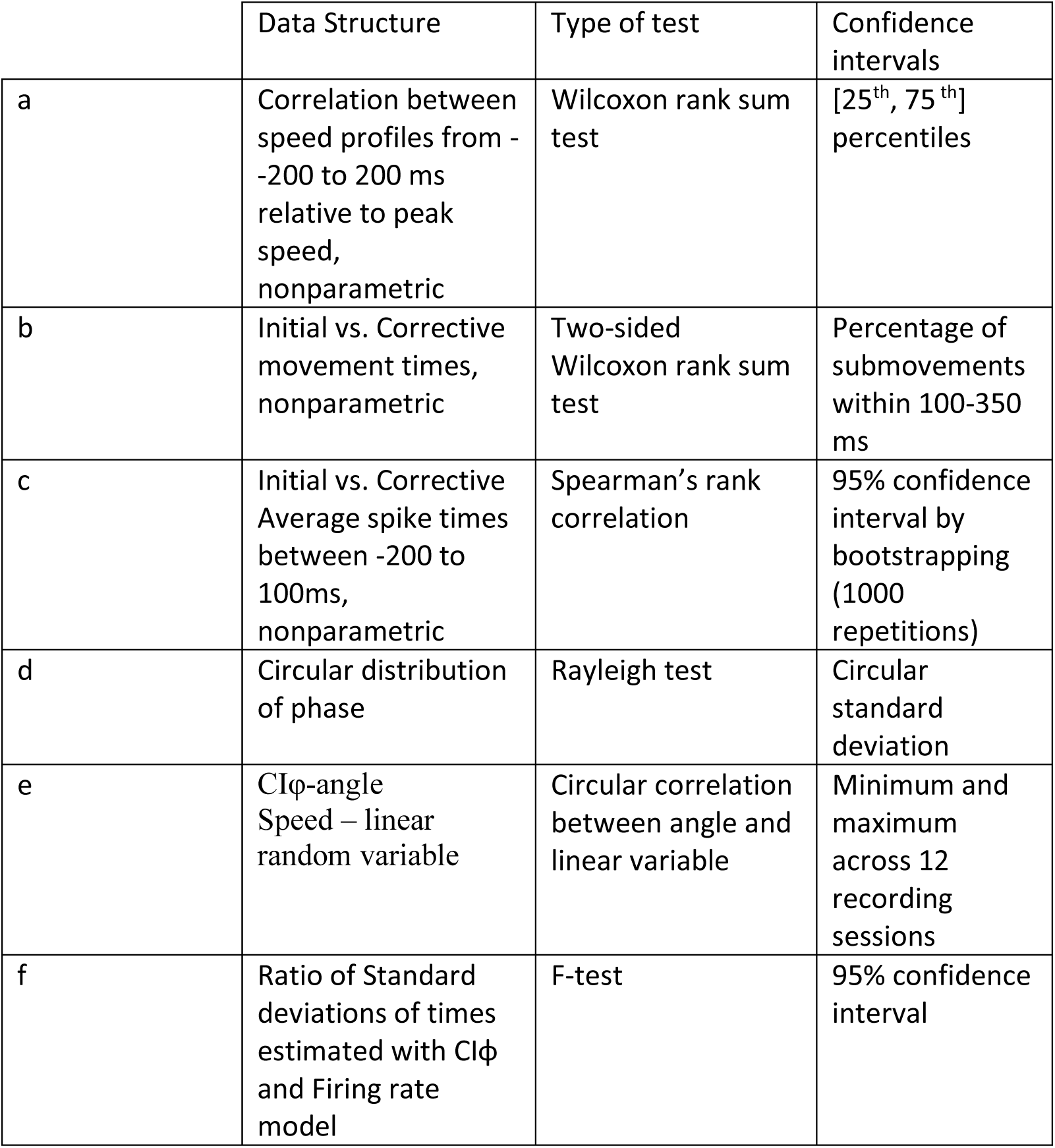
Statistical tests and confidence intervals reported throughout this study referenced with letter superscripts.

## Results

### Motor behavior – initial and corrective submovements

Movement speed was analyzed throughout the center-out task from instruction until successful completion of the final target hold. The two monkeys successfully completed 10,963 (monkey P) and 8,737 (monkey Q) trials across 12 recording sessions each. In addition to the peaks in speed with the initial reach after instruction, additional peaks in speed were observed and labeled as corrective submovements. There were 6478 and 3912 corrective submovements identified for monkeys P and Q, respectively. Across all trials, 68.3% (P) and 71.1% (Q) were completed in a single initial movement, 17.5% (P) and 20.3% (Q) of trials were completed with one additional corrective submovement, and 14.2% (P) and 8.6% (Q) of trials required two or more corrective submovements. The location of the identified speed peaks within example trials and the speed profiles for monkey P are shown in Figure 2A and 2B, respectively. The speed peaks tended to be distinct with nearly zero velocity between most peaks. As shown in Figure 2C, 99.0% (P) and 97.7% (Q) of the minimum speed trough following the initial speed peak were less than 20% of the peak. Similarly, 82.4% (P) and 85.8% (Q) of the troughs were less than 20% of the preceding peak between sequential corrective speed peaks. The mean peak speeds for initial submovements were 1533 (P) and 1182 (Q) pixels/s while corrective submovement peak speeds were 460 (P) and 400 (Q) pixels/s. Thus, the average peaks for corrective submovements were 30.0% and 33.8% of initial submovements, and a low-speed trough almost always occurred between two speed peaks making it reasonable to analyze submovements defined by their peak speeds.

**Figure 2.**
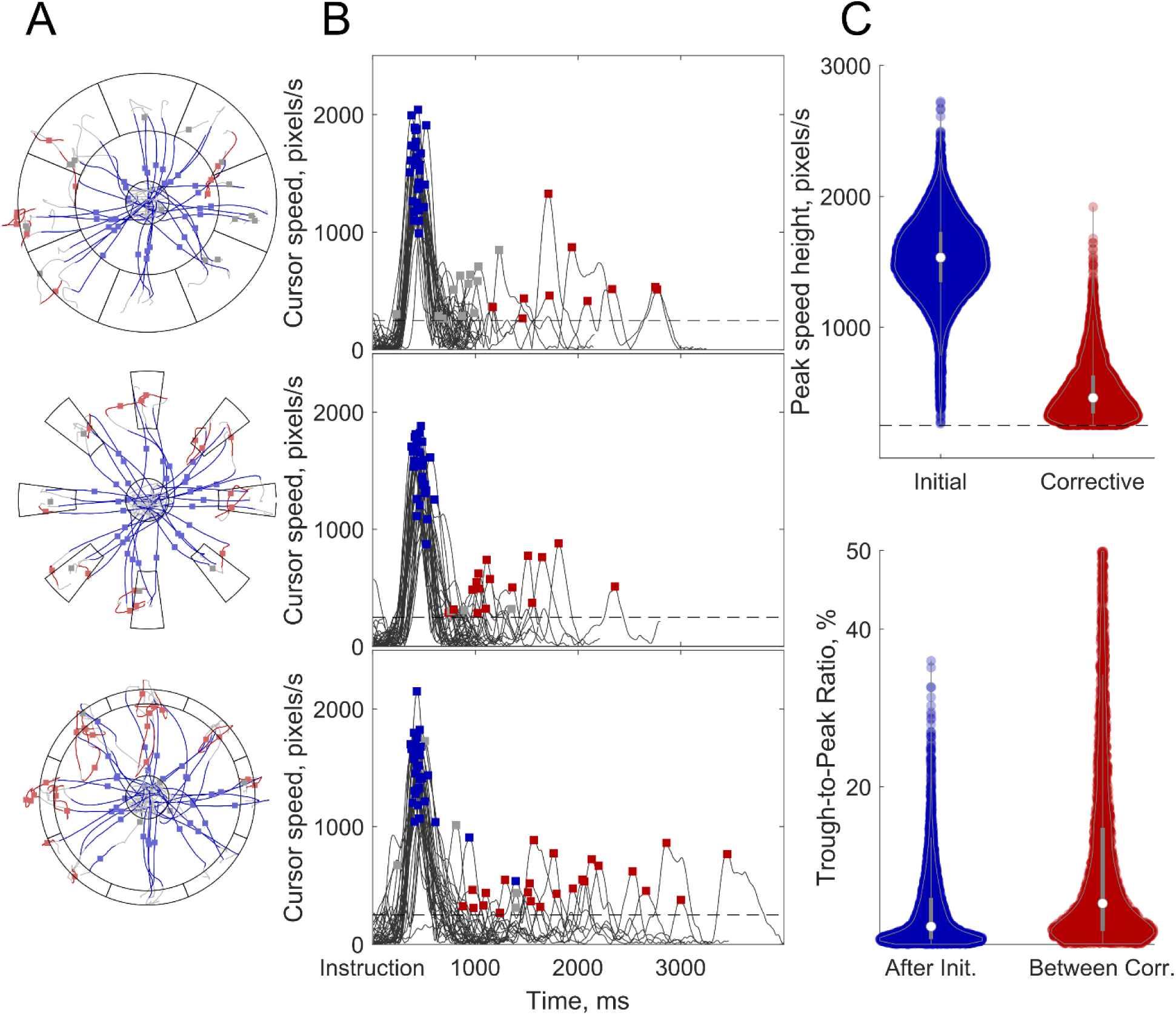
The precision center-out task. A) Cursor paths for four example trials to each target for the three target sizes: regular (top), narrow (middle), shallow (bottom). Initial submovements from 200ms before to 200ms after speed peaks are plotted in blue with the point when peak speed occurred shown with a blue dot. Corrective movements are similarly identified in red with a red dot. Grey lines connect the rest of a trial before, between, or after submovements with a speed peak. B) Cursor speed plotted versus time for a subset of trials. Initial (blue) and corrective (red) submovement speed peaks are identified with squares. Gray squares identify speed peaks that were thrown out because they i) were small initial movements that did not move outside the center or ii) occurred entirely within the peripheral target. C) Top) Distribution of peak speeds for initial (blue) and corrective (red) submovements. Bottom) Distribution of the trough-to-peak ratio for the troughs following an initial submovement before a corrective submovements and following a corrective submovement before another corrective submovement. Data is shown for monkey P. Data for monkey Q, which had similar results, is not shown.

The speed profiles were time aligned to peak speed to better examine the identified submovements (Figure 3A). Almost all submovements show a clear bell-shaped profile for both the initial and corrective movements. The similarity between initial and corrective speed profiles was assessed by using the correlation between randomly selected pairs of movements. For random pairs (irrespective of trial) of one initial and one corrective submovement, the median correlation was 0.78 [0.58 0.89] (monkey P) and 0.83 [0.70, 0.90] (monkey Q). Thus, the shape of corrective submovements was significantly correlated with the shape of initial submovements (p<0.001^a^). As a ceiling comparison, the correlation between randomly selected pairs of initial submovements was observed to be 0.93 [0.86 0.96] (P) and 0.91 [0.80, 0.96] (Q). Even though the shape of initial-corrective pairs was significantly less correlated than the initial-initial pairs, corrective submovements still had a similarity measure that was a large percentage—84% (0.78/0.93) and 91% (0.82/0.91) —of that observed for initial-initial pairs.

**Figure 3.**
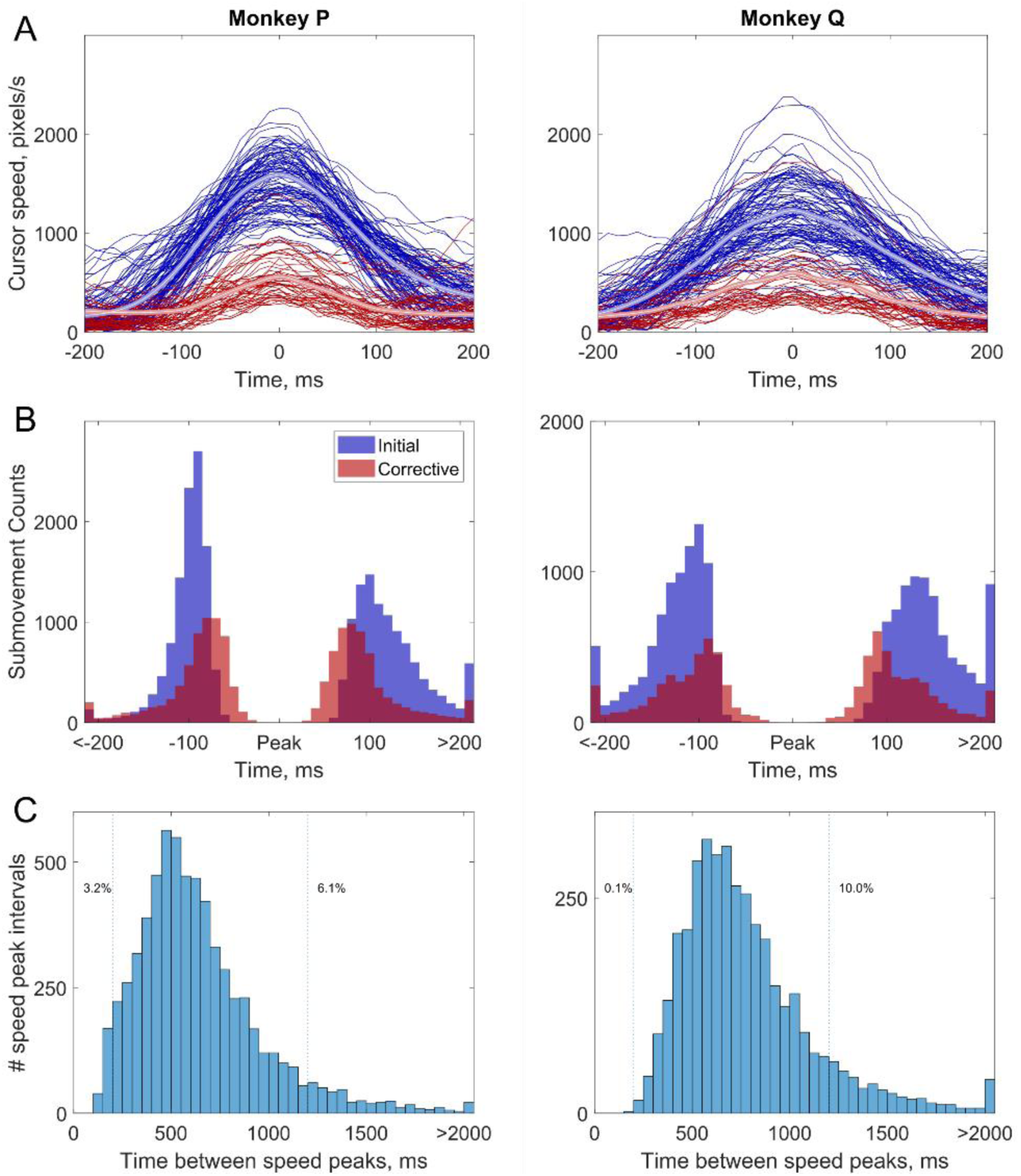
Time course of submovements. A) The cursor speeds are plotted aligned to speed peaks for initial (blue) and corrective (red) submovements. N.B. The cursor speeds shown are before the bandpass filter used for identifying peaks displayed in Figure 1B. Thus, the maximum of each trace may not align exactly with the plotted peak speed. B) Histogram of the time at half-maximum speed before and after peak speed for all initial (blue) and corrective (red) submovements. C) The time duration between speed peaks including the times from initial submovement to first corrective submovement as well as between any consecutive pairs of corrective submovements.

The time duration and timing of submovements was also examined. The onset and offset of submovements were defined as the time points when speed was one-half of the maximum speed both before and after the speed peak. As shown in Figure 3B, the movement duration at half maximum speed was similar and close to symmetric for both initial and corrective submovements. The initial submovements were slightly longer having a median time of 220ms (P) and 270ms (Q) compared to corrective submovements with medians of 180ms (P) and 220ms (Q). This difference in median movement times was statistically significant (p<0.001^b^) but the difference of 40 and 50ms was small, especially given the peak speed was only one-third the magnitude for the smaller corrective movements. Overall, all submovement durations, as measured by the full width at half maximum, occurred within a similar range with 96.7%/88.0% (P/Q) of all initial and 96.3%/93.0% (P/Q) corrective submovements between 100-350 ms. The time between speed peaks—either initial to first corrective submovements or between subsequent corrective submovements—is plotted in Figure 3C. The median time between peaks were 570 ms for monkey P and 700 ms for monkey Q with the mode time between peaks being 450ms (P) and 550ms (Q). Only 3.2% (P) and 0.1% (Q) of speed peaks had a time between peaks less than 200ms and 6.1% (P) and 10.0% (Q) of speed peak pairs had times greater than 1200 ms. These observations suggest the movement behavior could be divided into submovements with similar bell-shaped velocity profiles and similar time durations.

### Consistent Timing of Neural Firing Rates for Initial and Corrective Submovements

Single target acquisition movements thus often consisted of initial and corrective submovements with similar temporal characteristics. Did neural activity in the primary motor cortex control such target-acquisition movements as a single movement, or as a series of discrete submovements? The neural firing rates across the recorded population were time aligned to the submovement speed peaks to examine the firing rates from 500 ms before until 300 ms after the peak speed. The average firing rate (smoothed with a Gaussian window, σ=30ms) for all analyzed units aligned to the peak speed for initial and corrective submovements are shown in Figure 4A. A clear peak occurs before the peak speed for both initial and corrective submovements in both monkeys. Monkey P’s peak firing rates occurred 170 ms and 120 ms before initial and corrective submovements, respectively, while monkey Q’s occurred at 160 and 160 ms before for both initial and corrective submovements. Thus, firing rates increased and peaked globally for corrective submovements in addition to the initial reach.

**Figure 4.**
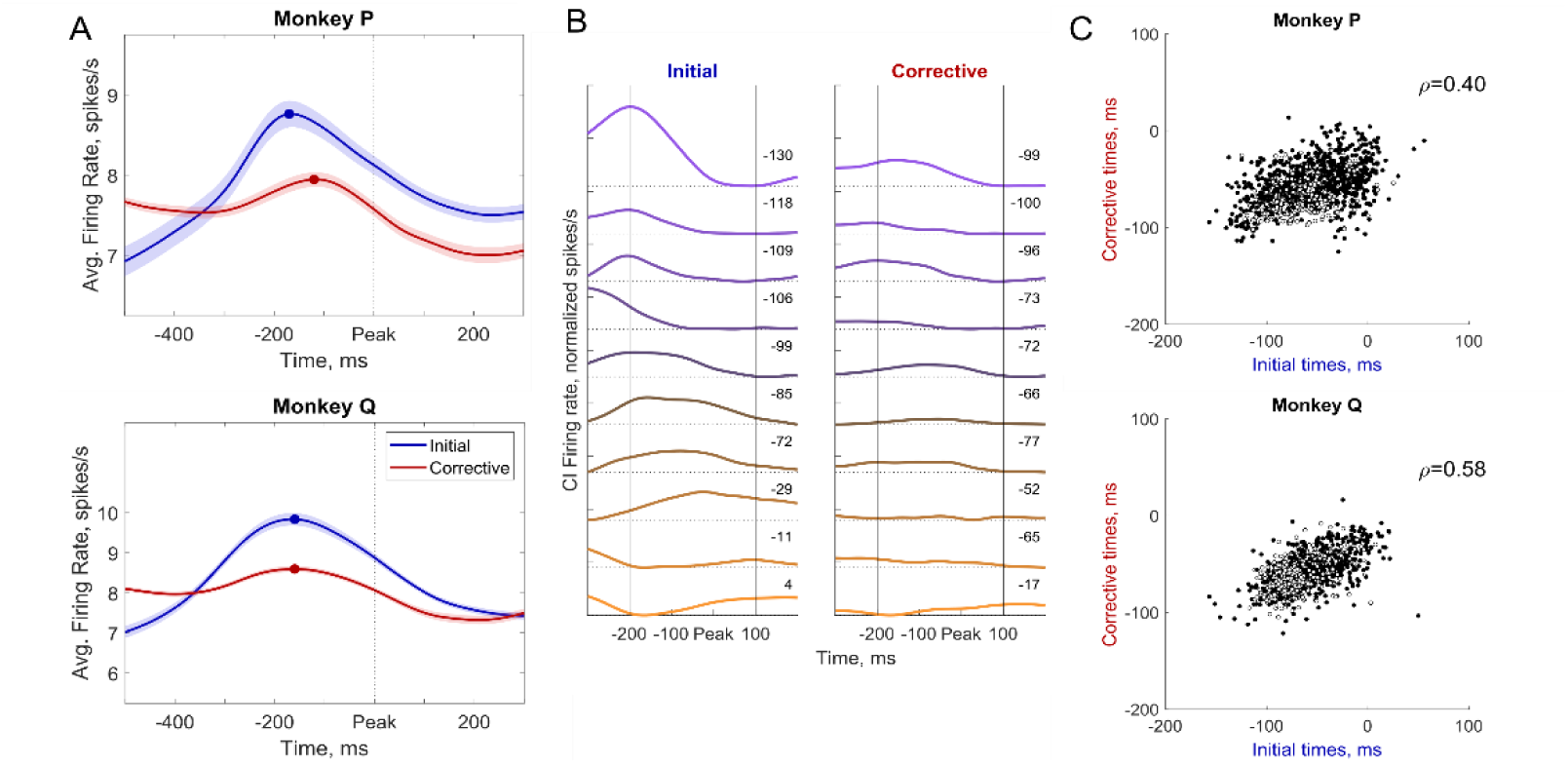
Neural firing relative to initial and corrective submovements. A) The firing rate for all spiking units was averaged for all initial (blue) and corrective (red) submovements. The shaded region interval shows the 95% confidence interval of the calculated mean for all spiking units. Circles indicate the time of peak firing rate for each condition. B) Average condition-independent firing rates for 10 example spiking units recorded simultaneously from monkey P time-aligned relative to peak speed for all initial (left) and corrective (right) submovements. Firing rates are shown relative to the average firing rate within the given time window (initial or corrective) for each spiking unit. The weighted timing of spikes (in ms) within the -200ms to 100ms window is given for each unit. Units are colored based on the initial movement by whether their firing rates were greater early (purple) or late (orange). C) Weighted timing of spiking relative to peak speed for each unit for initial (abscissa) and corrective (ordinate) submovements. More negative times represent spiking earlier relative to the peak speed of each submovement. Single units are shown with filled circles while all other spiking multi-units are shown with open circles.

If all neurons had the same time lag preceding the upcoming peak in movement speed, there would be a synchronized increase and decrease of all condition-independent firing rates simultaneously. However, when examining average firing rates from 10 example neurons from one recording session from monkey P, all aligned to peak speed, we see heterogenous timing of firing rates relative to the peak speed (Figure 4B). This relationship tended to be conserved across initial and corrective movements, with the purple spiking units tending to fire earlier and the orange units later for both initial and corrective submovements. This suggests that the condition-independent neural activity across the neurons might form a repeatable temporal structure—a neural trajectory—that is more than a simple simultaneous rise and fall in firing rate across the population

To quantify the early versus late consistency of spiking units, we calculated the average time of all spikes that occurred within a window from -200ms before to 100ms after peak speed to determine whether a unit tended to increase its firing rate earlier (negative time) or later (positive time) relative to peak speed. We then compared these average spike times for initial versus corrective submovements for each spiking unit. As shown in Figure 4C, earlier firing units (more negative) for initial submovements tended to fire earlier for corrective submovements, while units later (more positive) for initial submovements also tended to fire later for corrective submovements. This correlation was significant for all spiking units with Spearman correlations of ρ = 0.40 [0.35, 0.45] (P) and ρ = 0.58 [0.53, 0.62] (Q), p<0.001^c^. Using only single units, the Spearman correlations were ρ = 0.37 [0.31, 0.44] (P) and ρ = 0.61 [0.54, 0.68] (Q), p<0.001^c^. Thus, a significant portion of the ordered timing of units was conserved relative to peaks in movement speed for both initial and corrective submovements.

### Consistent Neural Dynamics for Initial and Corrective Submovements

We next wanted to examine whether these repeatable neural patterns that occurred on average across all movements could be used to identify submovements on individual trials. Despite the smaller magnitude of the condition-independent neural activity during corrective movements, the repeated oscillations in speed and repeated neural dynamics suggested a portion of neural activity was repeatable and common to initial and corrective submovements. To examine this, we built a simple linear dynamical system model using the neural firing rates from the entire trial—including both initial and corrective submovements—to characterize common temporal dynamics that might be present. The neural firing rates were again averaged across all conditions, i.e. movement directions, and both initial and corrective portions of the trials so the dynamical system model would identify common condition-independent activity. Using the jPCA algorithm described in Churchland et al. (2012), i) the first six principal components of the neural space and ii) the two dimension plane within the space of those six principal components that captured the most rotational neural activity were identified. We labeled the two neural dimensions of the plane with the most rotational condition-independent activity as CIx and CIy. To consistently define CIx and CIy across recording sessions and monkeys, we aligned the +CIx direction with the neural dimension that had the maximum average firing rate in the plane. This was performed by calculating the average firing rate across all spiking units for neural activity based on each timepoint’s angle in the CIx/CIy plane (binned in 100 angle intervals) and rotating the CIx and CIy axes so that +CIx aligned with the angle with largest firing rate. This alignment results in the +CIx dimension closely aligning with the time course of the global average firing rate across the population (shown in Fig 4A) while CIy is an orthogonal neural dimension that oscillates with a phase lag of π/2 compared to CIx.

The average firing rates projected in our identified CI plane for all initial and corrective submovements are shown in Figure 5, where the neural data was again aligned relative to peak speed for initial and corrective submovements separately. The neural trajectory in the 2-dimensional CIx/CIy plane are shown in Figure 5A, while the same CIx and CIy dimensions are plotted as a function of time in Figure 5B. The initial and corrective neural trajectories (Fig. 5A) are very similar in their shape and direction of rotation within the plane, with the trajectories for corrective submovements appearing as an additional cycle resembling a smaller, scaled version of the larger trajectories for initial submovements moving from the -CIy to +CIx to +CIy to -CIx dimensions. The time courses of CIx (solid) and CIy (dashed) (Fig. 5B) were similar for initial (blue) and corrective (red) submovements, though they differed in magnitude. The peak in the CIx dimension (denoted with an X)—defined as the dimension in the plane that best correlated with the global average firing rate of the population—occurred approximately 150 ms before peak speed for initial and corrective submovements, whereas the peak in the CIy dimension (also denoted with an X) occurred near the time of peak speed for both submovement types.

**Figure 5.**
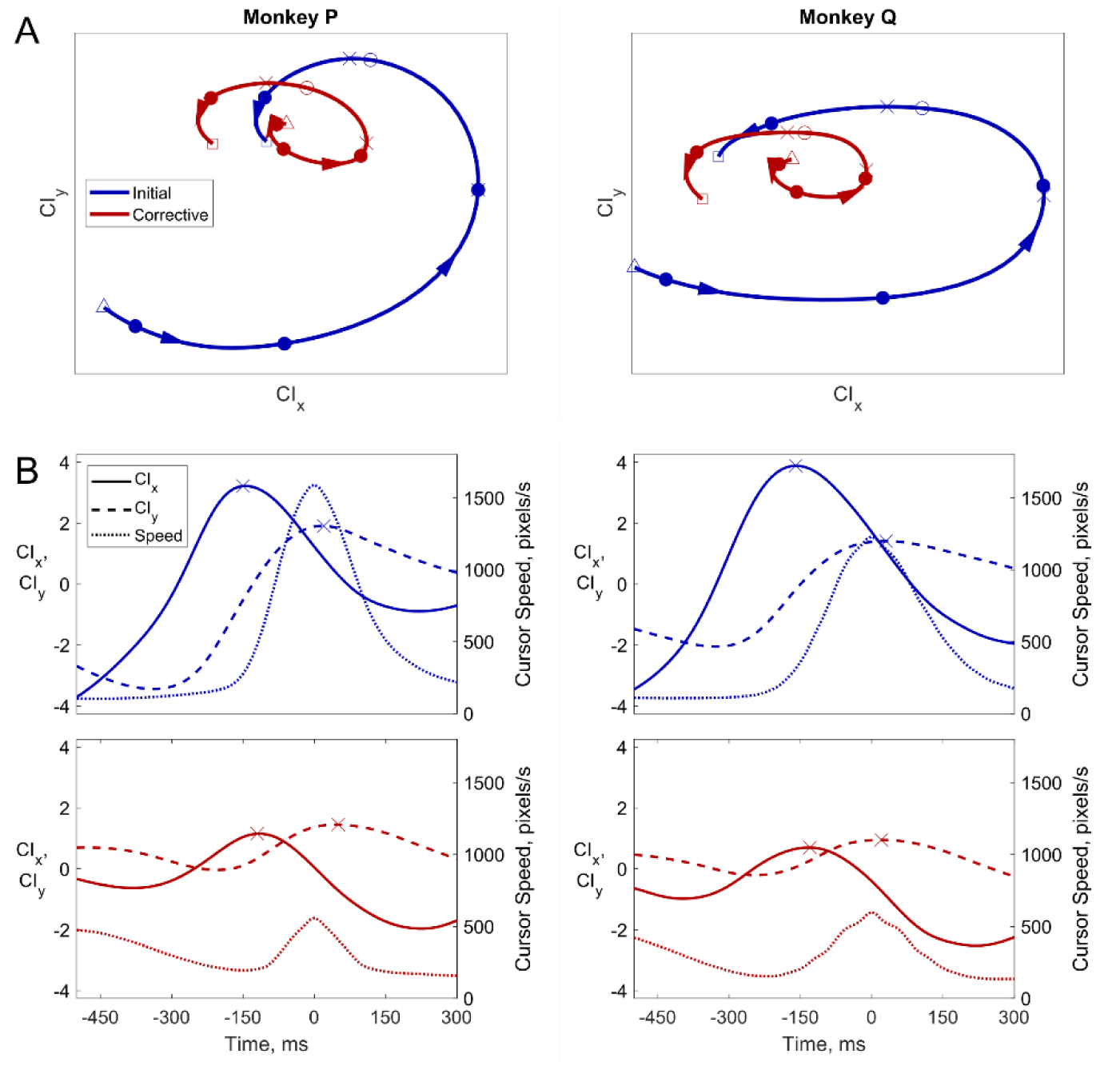
Cyclic neural dynamics related to initial and corrective submovements. A) The average population firing rates for initial (blue) and corrective (red) submovements are projected in the CIx/CIy plane identified with jPCA. The trajectories start at the triangles and end at the squares. Each filled circle is a 150 ms time step and the open corresponds to peak speed. C). Average CIx (solid lines) and CIy (dashed lines) plotted as a function of time relative to average cursor speed (dotted lines).

### Neural cycles improve predictions of behavioral timing

Since the population firing rates in the CI plane appeared to cycle across the two dimensions with similar timing for initial and corrective submovements, despite different magnitudes, we next chose to examine the instantaneous phases of CIx and CIy activity to see if it was a statistically significant marker of the neural state of motor cortex and its relationship with upcoming movement. We used a Hilbert transform to create an analytic representation of the CIx and CIy signals and then calculated the instantaneous phase by taking the angle between the real component and the Hilbert transformed imaginary component. The average phase of CIx and CIy for both initial and corrective submovements— time aligned to peak speed—is shown in Figure 6A. The phase of CIx (solid lines) and that of CIy (dashed lines) each were similar for initial and corrective submovements, with the zero phase of CIx occurring about 150 ms before the peak speed while CIy lagged CIx with an approximately π/2 phase lag, with the zero crossing occurring around peak speed. The slope of the phase for corrective movements was slightly steeper indicating that neural activity cycled slightly faster for corrective movements than initial. Histograms of the phase of CIx and of CIy at peak speed on individual trials are shown in Figure 6B. The distributions of phases of CIx and CIy were significantly non-uniform for both monkeys and the means and standard deviations are given in Table 2. Thus, there was a clear relationship between peak speed and the phase of condition-independent activity that occurred with almost all submovements, both initial and corrective, and had similar timing.

**Figure 6.**
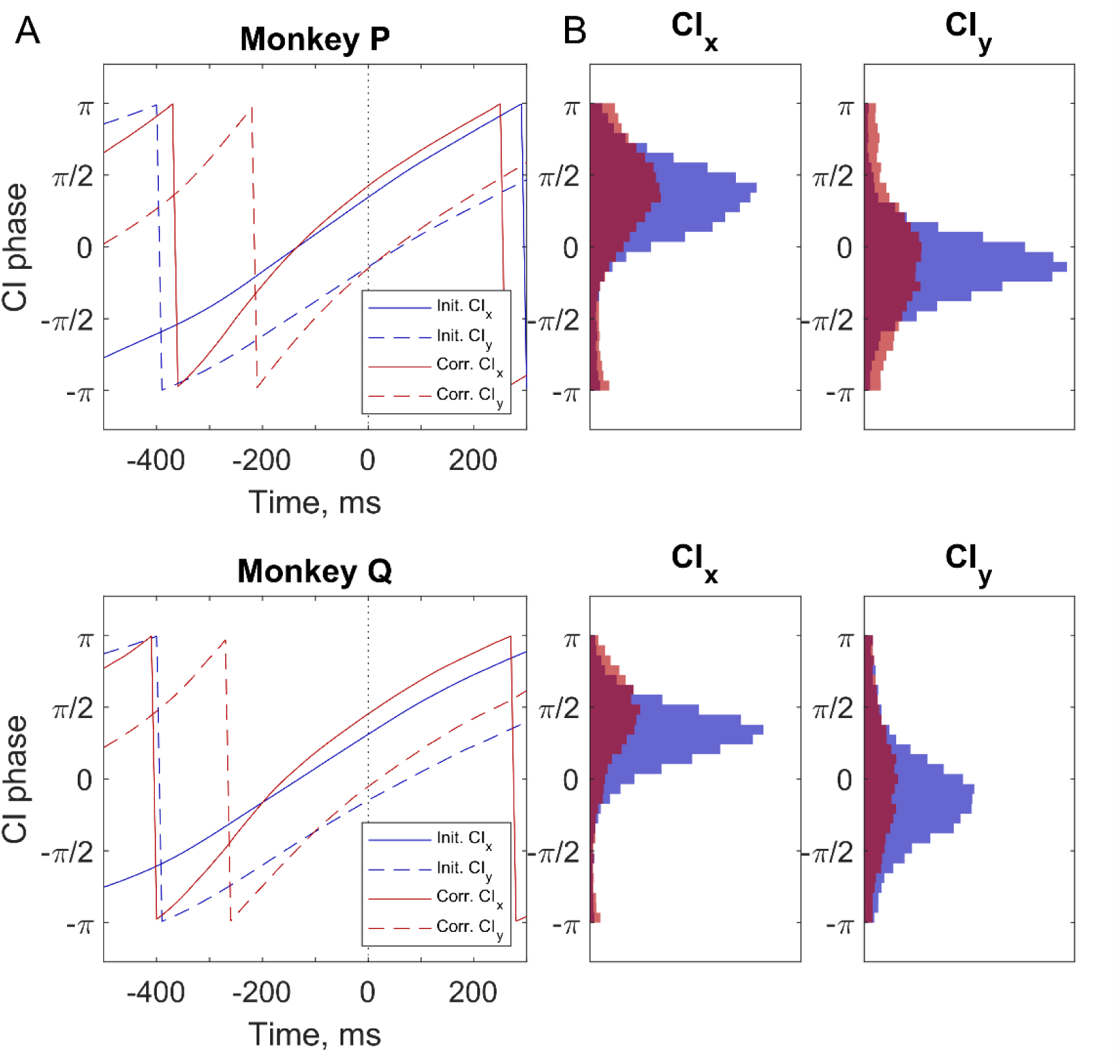
Phase of CIx and CIy relative to peak cursor speed. A) Phase of CIx (solid lines) and CIy (dashed lines) time-aligned to peak speed (Time = 0) and averaged for all initial (blue) and corrective (red) submovements. B) Histograms of the phase of CIx and Ciy at the time of peak speed for initial (blue) and corrective (red) submovements. Means and standard deviations are given in Table 2.

**Table 2.** Means and standard deviations of the phase of CIx and CIy. All circular distributions of the phase of CIx and CIy were non-uniform (all p < 0.001^d^).

| Monkey P | CIx Mean | CIx Std. dev | CIy Mean | CIy Std. dev |
| --- | --- | --- | --- | --- |
| Initial | $0.35 \pi$ | $0.26 \pi$ | $-0.14 \pi$ | $0.23 \pi$ |
| Corrective | $0.43 \pi$ | $0.33 \pi$ | $-0.15 \pi$ | $0.36 \pi$ |
| Monkey Q |  |  |  |  |
| Initial | $0.31 \pi$ | $0.22 \pi$ | $-0.15 \pi$ | $0.30 \pi$ |
| Corrective | $0.45 \pi$ | $0.31 \pi$ | $-0.05 \pi$ | $0.37 \pi$ |

Because the phase in the CI plane appeared to define the neural dynamics and predict upcoming speed peaks, we created a metric we call the condition-independent phase (CIφ) by averaging the phase of CIx and phase of CIy + π/2 to calculate the current phase in the CI plane. We then examined the continuous relationship between cursor speed and neural CIφ. In figure 7A, we have plotted the cursor speed as a function of CIφ. While the CIφ is an angle that ranges between +/-π radians when calculated, for purposes of display here we have incremented CIφ in steps of 2π to show how successive cycles of neural activity (abscissa) were related to movement speed (ordinate) as individual trials progressed through both initial and subsequent corrective submovements. The individual trials for monkey P in Figure 7A are the same as the trials shown in Figure 2B. However, the speed traces have now been stretched or condensed in time based on the current brain state measured with the CIφ. This plot now shows that the speed of movement varied with the cyclic neural activity with the cursor speeds for most trials rising and falling in 2π cycles of CIφ. Both the speed averaged across all trials (white) and the non-uniform occurrence of peak speeds in individual trials (black) demonstrate that movement speed was consistently correlated with the cycles of condition-independent neural activity. The statistically significant circular correlation between speed and CIφ was 0.44 [0.39,0.53] and 0.42 [0.35,0.50] (p<0.001^e^ for both animals) with the largest speeds occurring at CIφ = 0.32π and 0.31π (+2kπ) for monkeys P and Q, respectively.

**Figure 7.**
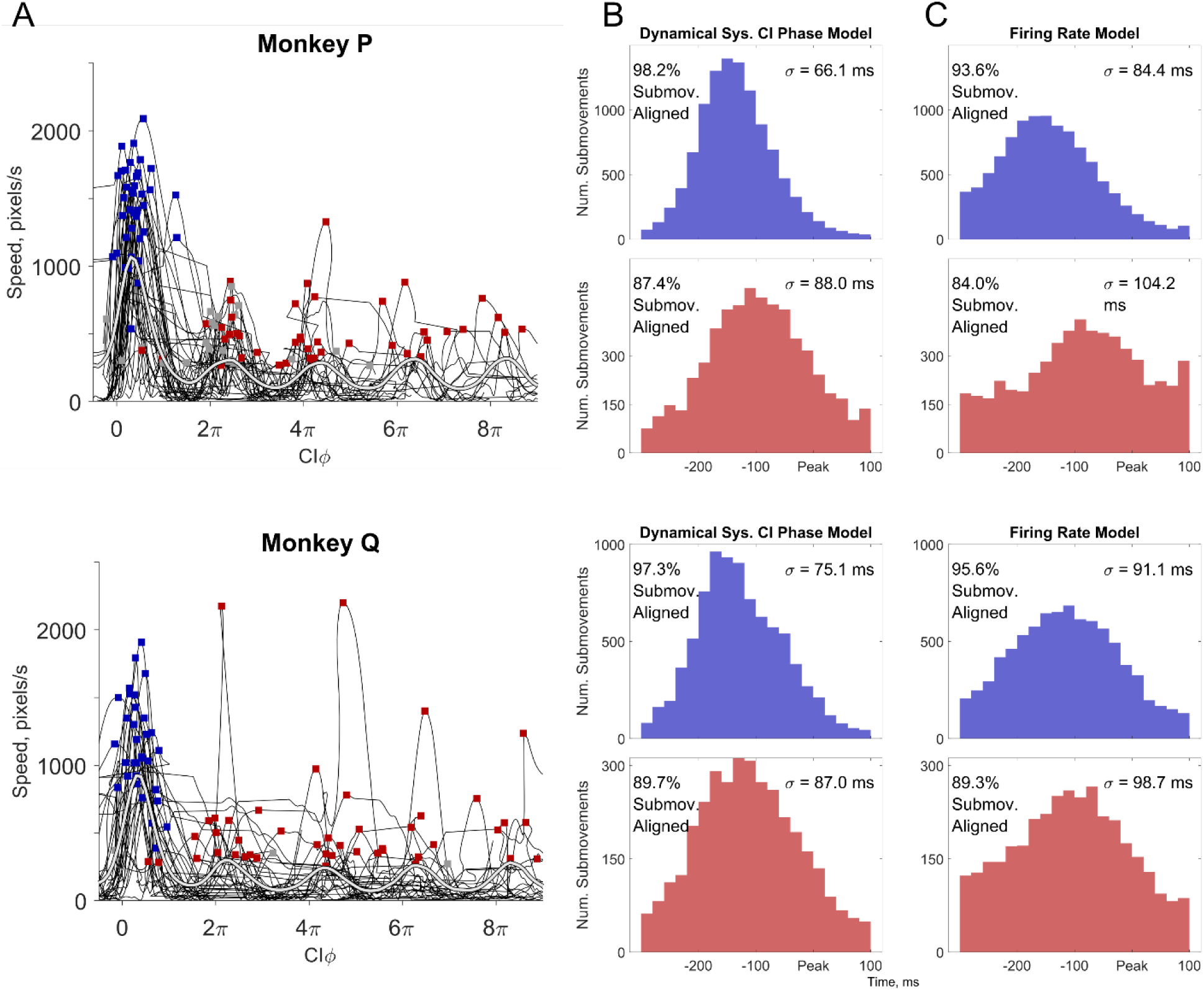
Relationship between CIφ and cursor speed. A) Cursor speed is plotted as a function of CIφ for 200 trials with at least one corrective submovement. The average speed of all trials as a function of CIφ is shown in white, illustrating the oscillation in cursor speed depending on the phase of neural activity.

Finally, we examined the predictive power of CIφ for estimating when the peak speed occurred. Figure 7B illustrates the distribution of the time at which CIφ =0 relative to the time of peak speed for initial submovements (top) and corrective submovements (bottom). These distributions consistently peaked 100-150 ms before the speed peak for both initial and corrective submovements. Corrective movements had CIφ =0 at times slightly closer to peak speed indicating that the time delay to peak speed was slightly less for corrective movements. A relatively consistent relationship between neural activity in the CIx/CIy plane and peak speed was present for both initial and corrective submovements across all trials regardless of target size or reach direction.

To examine if incorporating neural dynamics significantly improved prediction, we compared our CIφ predictions with these population dynamics to predictions using a standard approach of using the instantaneous firing rate of all units to predict peak speeds. For predictions with the instantaneous firing rates, we built a linear regression model to estimate speed with a weighted sum of the instantaneous firing rate (a single neural dimension) of all spiking units (see Methods). Using this model, we estimated the time when the peak in firing rate in the neural dimension occurred that predicted the upcoming speed peak. Figure 7C shows the temporal distributions of these peak firing rates relative to peak speed for both initial and corrective submovements. Like the distributions using the dynamical model above (Figure 7B), the firing rate model peaked 150 to 100 ms before peak speed. The peaks were broader by 10-20 ms, however, as characterized by the greater standard deviations (σ) given for each distribution.

The standard deviations were significantly different in all cases—initial and corrective for both monkeys (Table 2). Furthermore, although ≥ 84% of submovements were included in each of these distributions (percentages given in Fig. 7), a small fraction of submovements could not be aligned, lacking a CIφ=0 in the dynamical systems model and/or a peak in the firing rate model within the -300 to 100 ms time window examined. The percentage of these unaligned trials was consistently smaller for the dynamical systems model. Compared to using only the instantaneous/synchronous firing rates in a single neural dimension, using the cyclic/asynchronous dynamics of the neural population significantly improved the accuracy and consistency with which the time of peak speed could be predicted.

The circular correlations between CIφ and cursor speed for all corrective trials were 0.44 [0.39, 0.53] and 0.43 [0.36, 0.50] for monkeys P and Q, respectively, p<0.001^e^ in both cases. Note, the unwrapped CIφ is not always a monotonically increasing value as occasionally the neural activity could reverse and move clockwise rather than counter-clockwise in the neural plane shown in Figure 6B. B & C) Identifying the times of peak speeds with a dynamical systems model (B) or with an instantaneous firing rates (C). The time point when CIφ = 0 (B) or peak firing rate (C) was used as a prediction of the upcoming submovement. Each histogram shows only those submovements for which the neural data aligned with the movement data, i.e. CIφ = 0 (B) or maximum firing rate (C) occurred within the time range examined (−300 to 100 ms relative to submovement peak speed). The percentage of total aligned trials is shown for each distribution as well as the standard deviation (σ) for the aligned trials. In all cases, the dynamical systems model predictions were more precise, with a narrower standard deviation (statistics in Table 3) and fewer unaligned trials.

**Table 3.** Comparison of predication accuracy as measured with standard deviation in predictions using the dynamical system CIφ model vs. an instantaneous firing rate model.

| | $\sigma_1$,<br>CIϕ,<br>(ms) | $\sigma_2$,<br>Firing<br>Rate<br>(ms) | F-stat,<br>$\frac{\sigma_1^2}{\sigma_2^2}$ | 95%<br>Confid.<br>interval | Data<br>Comparison | Statistical<br>Test |
| --- | --- | --- | --- | --- | --- | --- |
| Initial |  |  |  |  | All<br>submovements<br>with a<br>prediction<br>between<br>-300:100 ms,<br>assuming<br>normal<br>distribution | F-test<sup>f</sup>, all<br>p<0.001 |
| Monkey<br>P | 66.1 | 84.4 | 0.61 | [0.59,<br>0.64] |  |  |
| Monkey<br>Q | 75.1 | 91.1 | 0.69 | [0.65,<br>0.71] |  |  |
| Corrective |  |  |  |  |  |  |
| Monkey<br>P | 88.0 | 104.2 | 0.71 | [0.68,<br>0.75] |  |  |
| Monkey<br>Q | 87.0 | 98.7 | 2.06 | [0.73,<br>0.83] |  |  |

## Discussion

Our precision center-out task utilized small targets to elicit one or more corrective submovements in many trials. We found a temporal relationship for both initial and corrective reaching movements with cyclic, condition-independent neural activity. Rather than a single cycle of neural activity in the primary motor cortex occurring during each trial, the speed profiles of initial and corrective submovements each aligned with a cycle of neural activity, providing a useful neural marker encoding the series of submovements.

In our precision center-out task, the monkeys’ movements showed consistent bell-shaped speed profiles. These speed profiles were evident for both the larger initial movement from the center toward the peripheral target as well as for each subsequent corrective movement. A large majority of both initial and corrective submovements had durations of 100-350 ms, with a low-speed trough separating almost all submovements. Discrete submovements defined by multiple speed peaks have previously been described in behavioral studies of reaching (Pratt et al., 1994; Lee et al., 1997; Hatsopoulos et al., 2007; Polyakov et al., 2009), turning a knob (Novak et al., 2000), isometric contractions (Massey et al., 1992; Hall et al., 2014), and object manipulation for tactile discrimination (Pruszynski et al., 2018). The experimental results and analysis presented here provide new evidence of a relationship between condition-independent neural dynamics and such behaviorally observed submovements.

### Condition-independent phase predictive of cursor speed

Churchland et al. (2012) originally described a single cycle of condition-dependent rotational dynamics in the activity of neurons in the primary motor and premotor cortex during both straight reaches and curved reaches around obstacles. More recently, Zimnik and Churchland (2021) demonstrated two repeated cycles of neural activity, each shortened in time, when a pair of movements were simultaneously instructed to be performed in rapid succession. Here, by focusing on the shifting dimensions of condition-independent neural activity with time, we identified that cycles of neural activity appear not only for initially planned reaches but also for the highly variable, corrective submovements that are made online with visual feedback. Our results highlight that the various time lags between individual cortical neurons’ firing and the upcoming reaching movements are conserved, whether large and instructed or small and made online with feedback.

### Similar but smaller cyclic, condition-independent activity for corrective movements

Although the orientation and direction of rotation through the identified condition-independent neural dimensions was similar for initial and corrective submovements, the magnitude of the condition-independent neural activity that occurred for corrective submovements was approximately one-third to one-half the magnitude of that for the initial submovements (both in average firing rate, Fig. 4A, and within our identified rotational CI plane, Fig. 5). On average, the encoding of movement speed is clearly present in primary motor cortex (Moran and Schwartz, 1999; Paninski et al., 2004), and the smaller change in average firing rate observed here during corrective movements reflected the lower movement speed for the corrective compared to the initial submovements, suggesting speed tuning in the magnitude of the condition-independent activity. This does not imply, however, that each individual trial and each individual neuron have proportionally smaller changes of firing rate during smaller amplitude movements. Examination of small, instructed movements has shown that a fraction of primary motor cortex neurons have similar firing rates for small, precise and for larger wrist movements while others are selective for only larger movements (Fromm and Evarts, 1981). We too observed similar large changes in firing rate on individual corrective submovements for certain neurons (data not shown). Only when averaging firing rates—time aligned to the peak movement speed or the decoded condition-independent phase—were the population differences in firing rate modulation between initial and corrective movements readily apparent. Precisely identifying encoded speed on a trial-by-trial basis with the neural activity remains challenging as there are often large changes of firing rates for individual neurons that are variable and idiosyncratic during any particular corrective submovement.

Our results highlight that condition-independent neural signals can evolve in time along with the neural dynamics that are related to task conditions. Adding condition-independent activity to condition-dependent activity has been suggested to make brain dynamics more robust to noise by increasing the differences in neural signals even when the muscle activation pattern at certain time points are very similar (Russo et al., 2018). In the context of precise, corrective movements, we speculate cyclic brain dynamics can be used to organize neural activity that creates distinct submovements with time-varying neural and musculoskeletal dynamics that are more reliable for motor control. Previous reports of neural activity defining submovements linked together have used the term movement fragments (Hatsopoulos et al., 2007). In the context of precise movements, we hypothesize that organizing movement into submovements or movement fragments might allow the control of particular submovements to have different encoding features, neural processing, or control policies, for instance, allowing the large initial movements to be larger amplitude and less precise while the corrective submovements are smaller and more precise. Further studies will be needed to understand the condition-dependent differences that accompany the condition-independent neural features presented here.

Though various time lags in different neurons seem likely to be present across many tasks, cyclic, condition-independent neural dynamics may not be similar for all upper extremity movements. For instance, whereas during combined reach-and-grasp movements cyclic condition-independent activity occurs along with more complex condition-dependent dynamics (Rouse and Schieber, 2018), during separate reaching movements and grasping movements condition-dependent activity was cyclic during reaching, but was more complex during grasping (Suresh et al., 2020). The neural signals in a given hemisphere for cyclic movements of the contra- and ipsilateral arms have also have been reported to be in orthogonal subspaces (Ames and Churchland, 2019). Cyclic neural activity may not be due only to intrinsic neural dynamics in M1, but also the result of sensorimotor feedback control and/or a cognitive strategy. With sufficient time delay between each submovement, the neural activity could fit both descriptions. Observations of additional submovements defined by second or third speed peaks do not necessarily require a feedback controller with discrete updates. A single, continuous optimal feedback controller with appropriate delays and signal dependent noise can generate additional submovements with multiple, sequential speed peaks (Li et al., 2018). Results by Susilaradeya et al. (2019) argue that extrinsic effects of a task interact with the intrinsic dynamics of the brain in a manner consistent with an optimal feedback controller, possibly providing a framework for assessing these effects across a variety of tasks including our precision center-out task. Further work examining neural activity in various tasks and/or additional sensorimotor brain areas will be needed to advance our understanding of the neural dynamics of the sensory processing, cognitive planning, and motor execution for precise, corrective movements.

The cyclic dynamics of corrective movements have important implications for brain-computer interfaces (BCIs). To date, most BCI decoders are time-invariant, not recognizing when submovements occur. Decoders are typically first constructed from observed or imagined movements that assume single, straight-line movements. When algorithms for updating BCI decoders consider the change in movement direction for corrective movements, it typically has been assumed the intended path is updated continuously (Gilja et al., 2012; Shanechi et al., 2016). Experiments have suggested that BCI control can be improved with two states: active control and rest (Kim et al., 2011; Williams et al., 2013, 2016; Sachs et al., 2016). Our results suggest that computing the phase of cyclic, condition-independent neural activity with CIφ (Fig. 7B) can provide better prediction of the timing of corrective submovements than using the instantaneous firing rates alone (Fig. 7C). This may lead to BCIs that allow the subject to better signal when they intend to make a corrective movement. With additional information about the typical neural dynamics and kinematics of submovements, BCI decoders may better estimate natural kinematics from noisy neural signals. Taking into account the cyclic dynamics of the condition-independent neural activity may also lead to better descriptions of the condition-dependent activity that encodes task features. For example, direction encoding has been shown to shift progressively during a single movement (Sergio and Kalaska, 1998; Churchland and Shenoy, 2007; Suminski et al., 2015; Suway et al., 2017). Accounting for the phase of a movement with its cyclic, condition-independent activity (i.e. CIφ) could enable decoders of movement direction that shift progressive during a single movement. Such improvements could lead to a more robust description of the neural encoding of precise and corrective movements.

Extended Data 1. Matlab code to calculate the CIφ is available on Github.

Since the trial data contains corrective movements in addition to the large initial movements that were not precisely time aligned to trial events for averaging condition-independent neural activity, we developed a novel algorithm to iteratively average the firing rates, calculate CIφ, then average the firing rates again based on the CIφ. This iterative process involves three steps: i) Each unit’s firing rate is averaged across all trials to determine its condition-independent firing rate. ii) Dimensionality reduction is performed using PCA and jPCA on the condition-independent firing rates to identify the neural plane with the most rotational/cyclic condition-independent activity. iii) The instantaneous phase is calculated using the Hilbert transform on the first two jPC dimensions for all data points. Matlab code is available on GitHub. Further details are available in the Readme document attached to the code.

## Acknowledgements

This work was supported by NIH NINDS K99/R00 NS101127.

